# Generation of promoters enabling high-level constitutive gene expression in both plants and *Escherichia coli*

**DOI:** 10.64898/2026.05.17.725692

**Authors:** Pabasara Rasadaree Weerasinghe, Daisuke Tsugama

**Affiliations:** Asian Research Center for Bioresource and Environmental Sciences (ARC-BRES), Graduate School of Agricultural and Life Sciences, The University of Tokyo, 1-1-1 Midori-cho, Nishi-tokyo-shi, Tokyo 188-0002, Japan

**Keywords:** Bacteria, Plant, Synthetic promoter, Selection marker

## Abstract

Functional validation of genetic components in plants often requires cloning them separately into both plant and bacterial expression vectors, a process that is both time-consuming and laborious. This study aimed to simplify this workflow by developing plant-bacteria dual-host promoter systems that drive high-level constitutive expression in both environments. To achieve this, two variants of the chloramphenicol acetyltransferase promoter (PCAT), a bacterial σ factor-dependent promoter, were integrated into the cauliflower mosaic virus 35S promoter (P35S), and their performance was evaluated using a hygromycin phosphotransferase (HPT)-GFP fusion reporter. One of these variants, PCAT1, conferred hygromycin resistance to *Escherichia coli* (DH5α and BL21 (DE3)) and maintained high-level expression comparable to the original P35S in onion epidermal cells. A hybrid P35S enhancer-PNOS system also conferred hygromycin resistance to *E. coli*, but its activity in inducing GFP signals in onion cells remained lower than that of P35S. Due to its compact size (89 bp) and efficiency, PCAT1 can serve as a module for converting standard plant vectors into dual-host systems, accelerating gene characterization and the development of new gene-based tools.

## Introduction

In plant molecular biology, the functional validation of genetic components such as selectable markers, reporter proteins and genome-editing tools is necessary for implementing bioengineering of interest. A bacterial expression system (most often an *Escherichia coli* expression system) enables rapid investigation of protein functions such as enzyme activity and protein-protein interactions and is therefore useful for characterization of any genes, even when they are derived from plants or used in plants. Conventionally, plant-bacteria dual-host evaluation necessitates the subcloning of a gene of interest into two separate expression vectors (i.e., a plant expression vector and a bacterial expression vector), which increases both the experimental workload and duration required. Streamlined “dual-expression” systems that function efficiently in both prokaryotic and eukaryotic environments can save such labor and time and thereby contribute to promoting gene characterization.

Achieving strong expression across plants and bacteria within a single regulatory unit remains a challenge. A common approach utilizes the T7 promoter and terminator. While these sequences are exceptionally compact (∼20 bp for the promoter and ∼50 bp for the terminator), their activities are dependent on the presence of T7 RNA polymerase (Studier and Moffatt, 1986), which limits suitable bacterial hosts to specific strains, such as λDE3 lysogens. Although broad-spectrum promoters functional across bacteria, yeast, and plant cells have recently been explored by stacking various regulatory elements (Morozov et al., 2025), achieving high-level constitutive expression specifically in bacterial hosts remains a challenge without relying on specialized machinery such as T7 RNA polymerase. Alternatively, the *Agrobacterium*-derived nopaline synthase (NOS) promoter (PNOS) and terminator (TNOS) are known to function in both plants and bacteria (e.g., Jefferson et al., 1987). However, the transcriptional activity of PNOS in plants is relatively weak and potentially insufficient for applications requiring high-level gene expression.

The present study describes the design of novel plant-bacterial promoter systems for achieving constitutive and high-level gene expression in both hosts. One system integrates a compact, bacterial sigma factor-dependent promoter, the chloramphenicol acetyltransferase promoter (PCAT) (e.g., Hogan et al., 2019), into the cauliflower mosaic virus 35S promoter (P35S) (Odell et al., 1985), which is widely utilized for its strong constitutive activity in plants. The PCAT sequence was selected for its minimal length and low frequency of internal ATG codons, which simplifies the removal of potential upstream open reading frames (uORFs) that could interfere with downstream translation in plant cells. The other system employs a hybrid approach, fusing the 35S enhancer region to PNOS to augment its expression levels in plant cells. These systems were validated using a *hygromycin phosphotransferase* (*HPT*)-*GFP* fusion gene as a reporter. We confirmed that these constructs can induce strong gene expression in both *E. coli* and onion epidermal cells, providing a versatile module for converting standard plant expression vectors into efficient plant-bacteria dual-host systems.

## Materials and Methods

### Bacterial strains and growth conditions

*Escherichia coli* strain DH5α was used for all cloning and plasmid construction. Cells were cultured in Luria-Bertani (LB) medium (10 g/L tryptone, 5 g/L yeast extract, and 10 g/L NaCl). Plasmid DNA was isolated using the GenElute Plasmid Miniprep Kit (Sigma-Aldrich, St. Louis, MO, USA) according to the manufacturer’s instructions.

### Construction of the base plant expression vector

To generate a streamlined plant-bacteria dual-expression backbone, pBI121-35SMCS-GFP (Tsugama et al., 2012) was digested with *Hind*III and *Eco*RI. The resulting fragment containing the P35S-GFP-TNOS cassette was gel-purified and ligated into the *Hind*III- and *Eco*RI-digested backbone of pER8 cHA-FG1 (Zuo et al., 2000; Pavicic et al., 2017; a gift from Kristiina Himanen (Addgene plasmid #118689; http://n2t.net/addgene:118689; RRID: Addgene_118689)). This step replaced the original T-DNA region with the P35S-GFP-TNOS cassette, yielding pER8d-35SMCS-GFP. Subsequently, the coding sequence (CDS) of *HPT* was amplified via PCR using pCAMBIA1300 (Hajdukiewicz et al., 1994; Abcam, Cambridge, UK) as a template and the primers listed in Supplementary Table S1. The *HPT* fragment was inserted into the *Sal*I-*Spe*I site of pER8d-35SMCS-GFP using the In-Fusion Snap Assembly Master Mix (Takara Bio, Kusatsu, Japan) to generate pER8d-35S-HPT-GFP. Transformants were selected on LB agar supplemented with 100 mg/L spectinomycin (Spec).

### Design and integration of the PCAT and PNOS

The PCAT sequence was derived from pAH25-SceI (Hogan et al., 2019; Addgene plasmid #129389). The start codon (ATG) was removed, and the sequence was manually truncated and modified to optimize its strength as a σ factor-dependent promoter, utilizing the iPro70-PseZNC predictor (Lin et al., 2019). The resulting optimized sequence was synthesized as two partially complementary oligonucleotides (Supplementary Table S1). These were annealed by heating at 95°C for 5 min followed by cooling to room temperature for 5 min, producing a double-stranded fragment with *Sal*I-compatible ends. This fragment was ligated into the *Sal*I-digested pER8d-35S-HPT-GFP and transformed into chemically competent DH5α cells prepared as previously described (Inoue et al., 1990). Selection was performed on LB agar containing 300 mg/L hygromycin B (Hyg) (Fujifilm Wako, Osaka, Japan) to screen for functional promoter activity. Two large colonies were selected for plasmid isolation and sequencing. One of the identified sequences contained mutations (designated as PCAT1), while the other was identical to the original design (designated as PCAT2).

For the PNOS constructs, the PNOS fragment was amplified from pBI121 (Jefferson et al., 1987) via PCR using the primers listed in Supplementary Table S1. The product was digested with *Sal*I and ligated into the *Eco*RV-*Sal*I sites of pER8d-35S-HPT-GFP. After transformation into DH5α and selection on Hyg-containing medium, the inserted sequences of the two largest colonies were sequenced, which confirmed that both clones contained the PNOS sequence.

The sequences of P35S-PCAT1-HPT-GFP-TNOS, P35S-PCAT2-HPT-GFP-TNOS, and P35S enhancer-PNOS-HPT-GFP-TNOS are available in Supplementary Fig. S1.

### Assessing Hyg resistance in *E. coli*

To evaluate the bacterial functionality of the designed promoters, the Hyg resistance conferred by each construct were assessed in two *E. coli* strains: DH5α and BL21 (DE3). For DH5α, 10 μL of chemically competent cells were transformed with 0.2 ng of plasmid DNA (1 μL of a 0.2 ng/μL stock). Following a 30-min incubation on ice and a 50-s heat shock at 42°C, the cells were recovered in 400 μL of liquid LB medium at 37°C for 1 h. From the resulting suspension, 200 μL aliquots were separately spread onto LB agar containing 100 mg/L Spec and LB agar containing 300 mg/L Hyg. Plates were incubated at 37°C for 16 h (Spec) and 20 h (Hyg), then stored at 4°C prior to colony counting.

For BL21 (DE3), 10 μL of Champion 21 chemically competent cells (SMOBIO Technology, Hsinchu, Taiwan) were transformed using 100 ng of plasmid DNA (2 μL of a 50 ng/μL stock) following the same protocol described above. Transformed cells were spread onto Spec and Hyg selection media and incubated at 37°C for 16 h for both antibiotics.

The ratio of Hyg-resistant (Hyg^*R*^) colonies to Spec-resistant (Spec^*R*^) colonies was calculated for each construct as a measure of the relative promoter strength.

### Observation and quantification of GFP fluorescence in *E. coli*

BL21 (DE3) transformants were cultured overnight in liquid LB medium supplemented with 100 mg/L Spec at 37°C until the OD_600_ reached approximately 4.0. The cultures were then normalized to an OD_600_ of 3.9 using fresh LB medium. A 1 mL aliquot of each normalized culture was centrifuged to obtain a cell pellet, and the supernatant was discarded. The tubes containing the pellets were placed on a SafeImager 2.0 Blue-Light Transilluminator (Invitrogen, Carlsbad, CA, USA). Fluorescence images were captured in a dark room using a smartphone camera (iPhone 16, Apple Inc., Cupertino, CA, USA) equipped with a 510 nm long-pass emission filter (BA510F; Olympus, Tokyo, Japan). The captured RGB images were decomposed into their respective color channels, red (R), green (G) and blue (B), using ImageJ software (National Institutes of Health). The mean gray value was measured at three representative points on each pellet in both the G and R channels. To normalize the data and account for variations in pellet thickness or lighting, the average G value was divided by the average R value for each sample. The resulting G/R ratio was defined as the relative GFP fluorescence intensity. This procedure was performed for three independent colonies per construct, which were treated as three biological replicates.

### Observation and quantification of GFP fluorescence in onion cells

The constructs with PCAT1, PCAT2 and PNOS as well as pER8d-35S-HPT-GFP were co-introduced with pBS-35SMCS-mCherry (Tsugama et al., 2013) into onion (*Allium cepa*) epidermal cells as previously described (Tsugama, 2026). Briefly, 500 ng of each construct was mixed with 7.5 μL of gold particles (0.6-μm diameter, 60 mg/mL; InBio Gold, Hurstbridge, Australia) and an equal volume of polyethylene glycol (PEG) solution (50% (w/v) PEG3350, 20 mM MgCl_2_). The suspension was incubated for 5 min at room temperature and centrifuged at 10,000 × *g* for a few seconds. The gold particles were washed once with 70% (v/v) ethanol and once with 99.5% (v/v) ethanol and resuspended in 15 μL of 99.5% ethanol. After 10 seconds of sonication, three shots with 5 μL each were delivered to the epidermal cells of an onion scale leaf. Three biological replicates were performed for each GFP construct. Following bombardment, the samples were incubated at 28°C for 16 h. Fluorescence images were captured using a BX51 microscope equipped with a DP73 CCD camera and cellSens software (Olympus, Tokyo, Japan). GFP images were acquired with a 1-s exposure and no neutral density (ND) filter, while mCherry images were acquired with a 1-s exposure using a U-25ND25 filter.

For quantitative analysis, individual cells with clear mCherry signals were selected. Using ImageJ (National Institutes of Health), cell outlines were manually traced with the Freehand Selection tool and refined using the Fit Spline function. The mean fluorescence intensity was measured within these regions of interest (ROIs) for both GFP and mCherry channels. To account for non-uniform background fluorescence, local background correction was performed by generating a 5-pixel-wide margin surrounding each cell ROI. The mean intensity of this background margin was subtracted from the mean intensity of the corresponding cell; negative values were assigned an intensity of zero. To normalize for variations in transformation efficiency and optical focus, the background-corrected GFP intensity was divided by the background-corrected mCherry intensity. This GFP/mCherry ratio was used as the measure of relative promoter strength. Data were pooled from at least 14 cells per construct.

## Results and Discussion

### Generation of constructs with plant-bacterial promoters

In preliminary experiments, the base vector pER8d-35S-HPT-GFP failed to confer hygromycin (Hyg) resistance to *E. coli* DH5α, a common cloning host. To overcome this, PNOS and PCAT were integrated into the vector using Hyg selection-coupled cloning, which successfully yielded Hyg^*R*^ colonies. Sequence analysis of two colonies each revealed that both PNOS clones and one PCAT clone (PCAT2) contained the expected sequences. In contrast, the other PCAT clone (PCAT1) harbored a 5-bp deletion (‘CGGGC’) and a G-to-A substitution, resulting in an additional 9-bp upstream uORF (‘ATGAGCTAA’) compared to the original design (Fig. 1A). These three constructs, along with the original pER8d-35S-HPT-GFP, were used for subsequent functional analyses (Fig. 1B).

**Figure 1.**
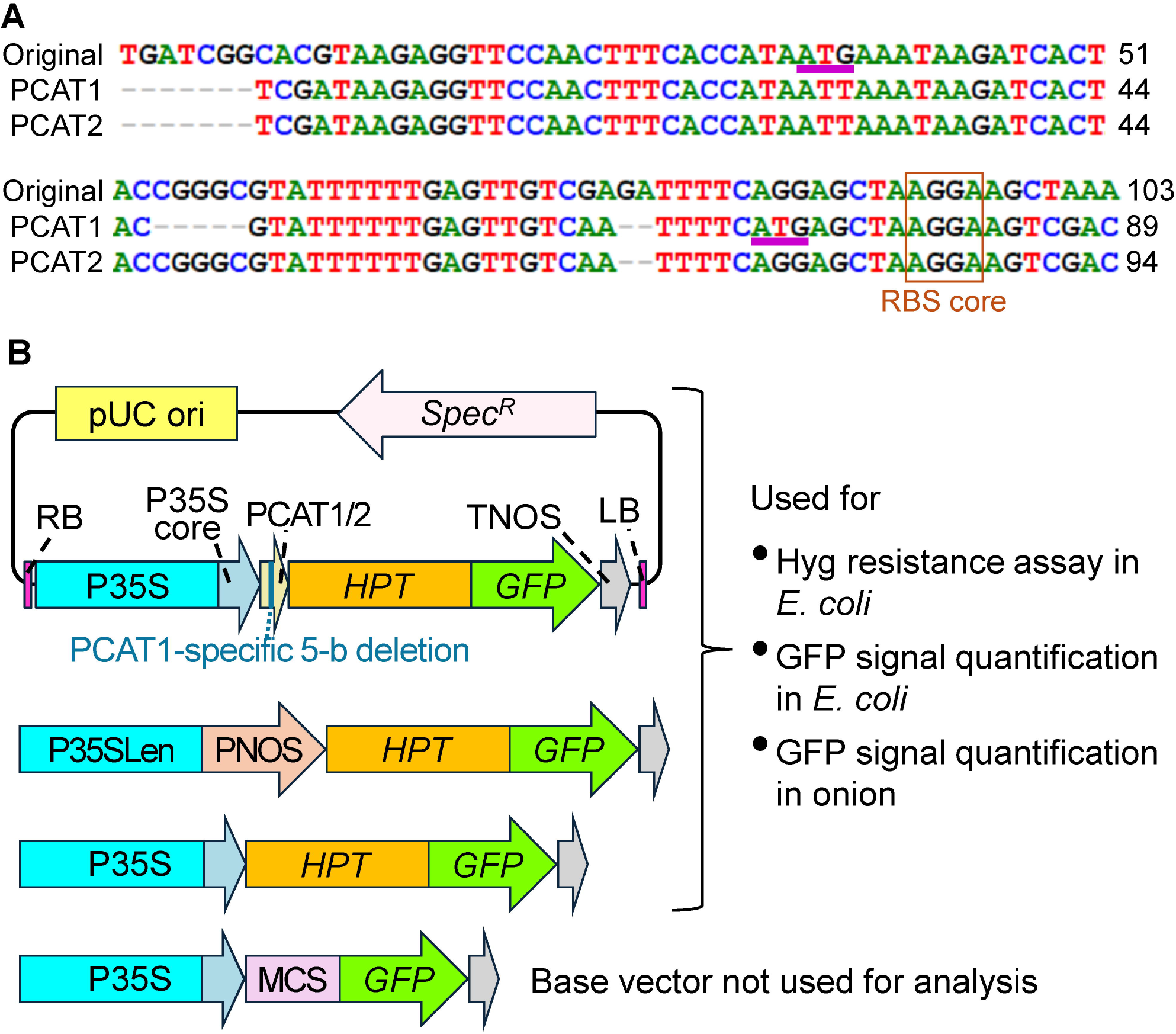
Constructs generated in this study. (A) Alignment of the PCAT variants. ‘Original’ refers to a commonly used PCAT variant, whereas PCAT1 and PCAT2 are two variants designed and cloned in this study. Numbers on the right indicate nucleotide positions. ATG, which can be used as the translation initiation site in eukaryotic cells, is underlined. AGGA proximal to the start codon represents the core of the ribosome-binding site (RBS) and is boxed. (B) The generated constructs and their applications in this study. P35SLen: P35S long enhancer, MCS: multiple cloning site.

### Performance of the constructs with plant-bacterial promoters in *E. coli* and plant cells

The PCAT1 and PNOS constructs conferred Hyg resistance to DH5α, whereas the PCAT2 construct and the control, base vector pER8d-35S-HPT-GFP did not (Fig. 2A, top panel). The failure of PCAT2 to confer resistance in this assay was unexpected, given its initial recovery from a Hyg^*R*^ colony. This discrepancy might be attributed to an undetected mutation outside the sequenced region or a growth-phase-dependent attenuation of resistance in DH5α. Nevertheless, all constructs except the base vector conferred Hyg resistance to BL21 (DE3), a strain optimized for protein expression (Fig. 2A, bottom; Fig. 2B). The PCAT2 and PNOS constructs induced stronger GFP signals than the base vector in BL21 (DE3) (Fig. 2C). These results suggest that the PCAT variants and PNOS can induce constitutive gene expression in *E. coli*, even when positioned downstream of P35S sequences.

**Figure 2.**
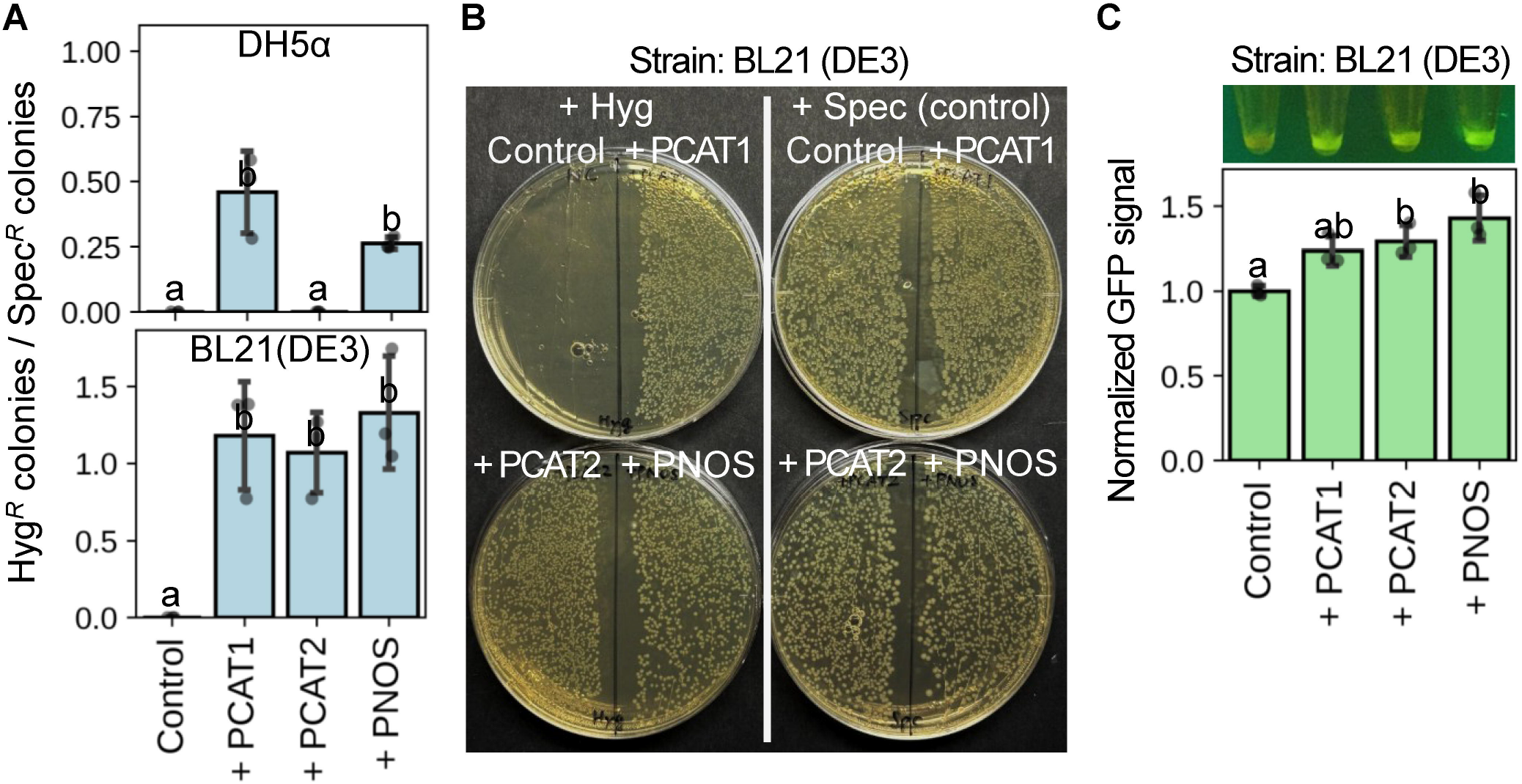
Activity of the newly generated promoters in *E. coli*. (A) Ratios of Hyg^*R*^ to Spec^*R*^ colonies obtained following transformation of *E. coli* strains DH5α (top panel) and BL21 (DE3) (bottom) with the indicated constructs. pER8d-35S-HPT-GFP was used as a control. Data are means ± SD from three replicates. Different letters indicate significant differences in a Tukey’s honestly significant difference test (*P* < 0.05). (B) Representative image of Hyg^*R*^ and Spec^*R*^ colonies for BL21 (DE3) transformants. (C) Relative GFP fluorescence intensities normalized to the red channel. Data are means ± SD from three replicates. Different letters indicate significant differences according to a Tukey’s honestly significant difference (HSD) test (*P* < 0.05).

In onion epidermal cells, the PNOS construct exhibited weaker GFP signals compared to the base vector. However, both PCAT1 and PCAT2 maintained GFP expression levels comparable to the original P35S (Fig. 3). These results suggest that while the 35S enhancer is insufficient to enhance PNOS activity to P35S levels, the integration of PCAT variants does not interfere with the high-level constitutive activity of P35S in plant cells.

**Figure 3.**
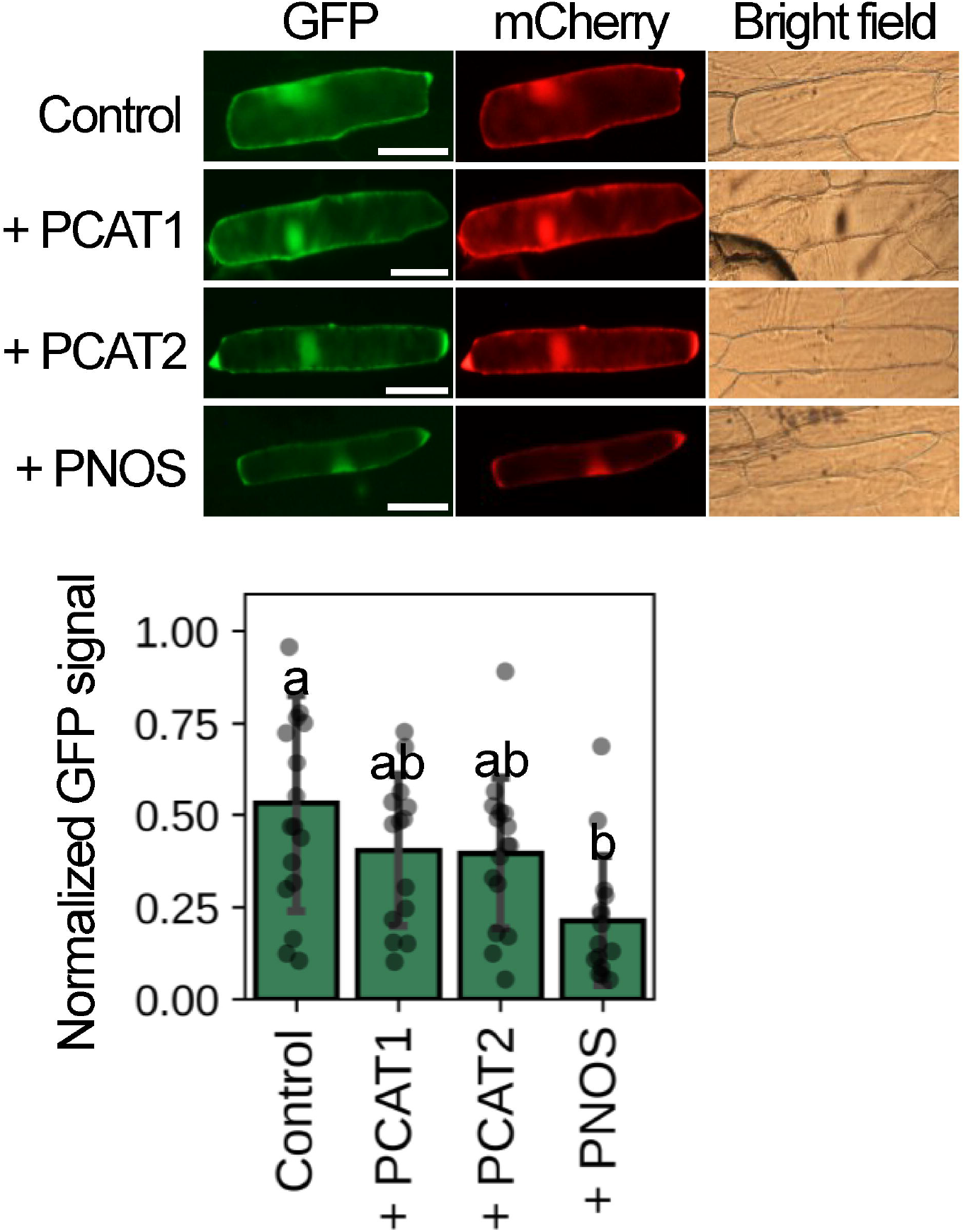
Activity of the newly generated promoters in onion epidermal cells. The control (pER8d-35S-HPT-GFP), PCAT1, PCAT2 and PNOS constructs were co-introduced with an mCherry-expressing construct into onion epidermal cells via particle bombardment. GFP and mCherry signals were detected 16 h post-bombardment. Top: Representative fluorescence images; scale bars = 100 μm. Bottom: Quantification of GFP signals normalized to mCherry signals. Data are means ± SD from at least 14 cells per construct. Different letters indicate significant differences according to Tukey’s honestly significant difference (HSD) test (*P* < 0.05).

Our data indicate that standard plant promoters can be effectively converted into plant-bacterial dual-host promoters by integrating PCAT variants or PNOS. Among these, PCAT1 is particularly promising; its compact size (89 bp, including the bacterial ribosome-binding site) allows for a simple design without compromising expression efficiency in either host.

## Abbreviations

CDS: Coding sequence
GFP: Green fluorescent protein
HPT: hygromycin phosphotransferase
Hyg: hygromycin
Hyg^*R*^: hygromycin-resistant
LB: Luria-Bertani
ND: neutral density
P35S: cauliflower mosaic virus 35S promoter
PCAT: chloramphenicol acetyltransferase promoter
PCR: polymerase chain reaction
PEG: polyethylene glycol
PNOS: nopaline synthase promoter
SD: Standard deviation
Spec: spectinomycin
Spec^*R*^: spectinomycin-resistant
TNOS: nopaline synthase terminator
uORF: upstream open reading frame

## Acknowledgments

The authors are grateful to Dr. Kristiina Himanen for providing pER8 cHA-FG1.

## Author contributions

PW and DT conceived the study and designed the experiments. PW and DT performed the experiments and data analysis. Both authors contributed to writing and revising the manuscript.

## Description of supplementary file

- Supplementary_Table_PCAT_PNOS.xlsx: Supplementary Table S1, which lists the primers and oligonucleotides used for this study
- Supplementary_Fig_S1_PCAT_PNOS.pdf: Supplementary Fig. S1, which is the sequences of P35S-PCAT1-HPT-GFP-TNOS, P35S-PCAT2-HPT-GFP-TNOS, and P35S enhancer-PNOS-HPT-GFP-TNOS.

## Data availability

All data generated or analyzed during this study are included in this published article and its Supplementary Information files.

