## Supplementary Fig. S1 for "Generation of promoters enabling high-level constitutive gene expression in both plants and *Escherichia coli*"

P35SLen-PNOS-HPT-GFP-TNOS

35S promoter long enhancer; modified NOS promoter (**putative RBS core**); HPT CDS; GFP CDS; NOS terminator

AGATTAGCCTTTTCAATTTCAGAAAGAATGCTAACCCACAGATGGTTAGAGAGGCTTACGCAGCAGG
TCTCATCAAGACGATCTACCCGAGCAATAATCTCCAGGAAATCAAATACCTTCCCAAGAAGGTTAAA
GATGCAGTCAAAAGATTCAGGACTAACTGCATCAAGAACACAGAGAAAGATATATTTCTCAAGATCA
GAAGTACTATTCCAGTATGGACGATTCAAGGCTTGCTTCACAAACCAAGGCAAGTAATAGAGATTGG
AGTCTCTAAAAAGGTAGTTCCCACTGAATCAAAGGCCATGGAGTCAAAGATTCAAATAGAGGACCTA
ACAGAACTCGCCGTAAAGACTGGCGAACAGTTCATACAGAGTCTCTTACGACTCAATGACAAGAAGA
AAATCTTCGTCAACATGGTGGAGCACGACACACTTGTCTACTCCAAAAATATCAAAGATACAGTCTC
AGAAGACCAAAGGGCAATTGAGACTTTTCAACAAAGGGTAATATCCGGAAACCTCCTCGGATTCCAT
TGCCCAGCTATCTGTCACTTTATTGTGAAGATAGTGGAAAAGGAAGGTGGCTCCTACAAATGCCATC
ATTGCGATAAAGGAAAGGCCATCGTTGAAGATGCCTCTGCCGACAGTGGTCCCAAAGATGGACCCC
CACCCACGAGGAGCATCGTGGAAAAAGAAGACGTTCCAACCACGTCTTCAAAGCAAGTGGATTGAT
GTGATATCATGAGCGGAGAATTAAGGGAGTCACGTTATGACCCCCGCCGATGACGCGGACAAGCC
GTTTTACGTTTGGAACTGACAGAACCGCAACGTTGAAGGAGCCACTCAGCCGCGGGTTTCTGGAGT
TTAATGAGCTAAGCACATACGTCAGAAACCATTATTGCGCGTTCAAAAGTCGCCTAAGGTCACTATC
AGCTAGCAAATATTTCTTGTCAAAAATGCTCCACTGACGTTCCATAAATTCCCCTCGGTATCCAATT
AGAGTCTCATATTCACTCTCAATCCAAATAATCTGCACCGGATC**AGGA**GGTCGACATGAAAAAGCCT
GAACTCACCGCGACGTCTGTCGAGAAGTTTCTGATCGAAAAGTTCGACAGCGTCTCCGACCTGATG
CAGCTCTCGGAGGGCGAAGAATCTCGTGCTTTCAGCTTCGATGTAGGAGGGCGTGGATATGTCCT
GCGGGTAAATAGCTGCGCCGATGGTTTCTACAAAGATCGTTATGTTTATCGGCACTTTGCATCGGC
CGCGCTCCCGATTCCGGAAGTGCTTGACATTGGGGAGTTTAGCGAGAGCCTGACCTATTGCATCTC
CCGCCGTGCACAGGGTGTCACGTTGCAAGACCTGCCTGAAACCGAACTGCCCGCTGTTCTACAACC
GGTCGCGGAGGCTATGGATGCGATCGCTGCGGCCGATCTTAGCCAGACGAGCGGGTTCGGCCCAT
TCGGACCGCAAGGAATCGGTCAATACACTACATGGCGTGATTTCATATGCGCGATTGCTGATCCCC
ATGTGTATCACTGGCAAACTGTGATGGACGACACCGTCAGTGCGTCCGTCGCGCAGGCTCTCGATG
AGCTGATGCTTTGGGCCGAGGACTGCCCCGAAGTCCGGCACCTCGTGCACGCGGATTTCGGCTCC
AACAATGTCCTGACGGACAATGGCCGCATAACAGCGGTCATTGACTGGAGCGAGGCGATGTTCGGG
GATTCCCAATACGAGGTCGCCAACATCTTCTTCTGGAGGCCGTGGTTGGCTTGTATGGAGCAGCAG
ACGCGCTACTTCGAGCGGAGGCATCCGGAGCTTGCAGGATCGCCACGACTCCGGGCGTATATGCTC
CGCATTGGTCTTGACCAACTCTATCAGAGCTTGGTTGACGGCAATTTCGATGATGCAGCTTGGGCG
CAGGGTCGATGCGACGCAATCGTCCGATCCGGAGCCGGGACTGTCGGGCGTACACAAATCGCCCG
CAGAAGCGCGGCCGTCTGGACCGATGGCTGTGTAGAAGTACTCGCCGATAGTGGAAACCGACGCCC
CAGCACTCGTCCGAGGGCAAAGAAAACTAGTATGGTGAGCAAGGGCGAGGAGCTGTTCACCGGGG
TGGTGCCCATCCTGGTCGAGCTGGACGGCGACGTAAACGGCCACAAGTTCAGCGTGTCCGGCGAG
GGCGAGGGCGATGCCACCTACGGCAAGCTGACCCTGAAGTTCATCTGCACCACCGGCAAGCTGCCC
GTGCCCTGGCCCACCCTCGTGACCACCTTCACCTACGGCGTGCAGTGCTTCAGCCGCTACCCCGAC
CACATGAAGCAGCACGACTTCTTCAAGTCCGCCATGCCCGAAGGCTACGTCCAGGAGCGCACCATC
TTCTTCAAGGACGACGGCAACTACAAGACCCGCGCCGAGGTGAAGTTCGAGGGCGACACCCTGGT

GAACCGCATCGAGCTGAAGGGCATCGACTTCAAGGAGGACGGCAACATCCTGGGGCACAAGCTGG
AGTACAACTACAACAGCCACAACGTCTATATCATGGCCGACAAGCAGAAGAACGGCATCAAGGTGAA
CTTCAAGATCCGCCACAACATCGAGGACGGCAGCGTGCAGCTCGCCGACCACTACCAGCAGAACAC
CCCCATCGGCGACGGCCCCGTGCTGCTGCCCGACAACCACTACCTGAGCACCCAGTCCGCCCTGAG
CAAAGACCCCAACGAGAAGCGCGATCACATGGTCCTGCTGGAGTTCGTGACCGCCGCCGGGATCAC
TCACGGCATGGACGAGCTGTACAAGTAAGAGCTCGAATTTCCCCGATCGTTCAAACATTTGGCAATA
AAGTTTCTTAAGATTGAATCCTGTTGCCGGTCTTGCGATGATTATCATATAATTTCTGTTGAATTAC
GTTAAGCATGTAATAATTAACATGTAATGCATGACGTTATTTATGAGATGGGTTTTTATGATTAGAG
TCCCGCAATTATACATTTAATACGCGATAGAAAACAAAATATAGCGCGCAAACTAGGATAAATTATCG
CGCGCGGTGTCATCTATGTTACTAGATCGGGAATTC

P35S-PCAT-HPT-GFP-TNOS

35S promoter long enhancer; modified PCAT (missing 5 b in the clone 1; [T or G in the clone 1 or clone 2, respectively]; **putative RBS core**); HPT CDS; GFP CDS; NOS terminator

AGATTAGCCTTTTCAATTTCAGAAAGAATGCTAACCCACAGATGGTTAGAGAGGCTTACGCAGCAGG
TCTCATCAAGACGATCTACCCGAGCAATAATCTCCAGGAAATCAAATACCTTCCCAAGAAGGTTAAA
GATGCAGTCAAAAGATTCAGGACTAACTGCATCAAGAACACAGAGAAAGATATATTTCTCAAGATCA
GAAGTACTATTCCAGTATGGACGATTCAAGGCTTGCTTCACAAACCAAGGCAAGTAATAGAGATTGG
AGTCTCTAAAAAGGTAGTTCCCACTGAATCAAAGGCCATGGAGTCAAAGATTCAAATAGAGGACCTA
ACAGAACTCGCCGTAAAGACTGGCGAACAGTTCATACAGAGTCTCTTACGACTCAATGACAAGAAGA
AAATCTTCGTCAACATGGTGGAGCACGACACACTTGTCTACTCCAAAAATATCAAAGATACAGTCTC
AGAAGACCAAAGGGCAATTGAGACTTTTCAACAAAGGGTAATATCCGGAAACCTCCTCGGATTCCAT
TGCCCAGCTATCTGTCACTTTATTGTGAAGATAGTGGAAAAGGAAGGTGGCTCCTACAAATGCCATC
ATTGCGATAAAGGAAAGGCCATCGTTGAAGATGCCTCTGCCGACAGTGGTCCCAAAGATGGACCCC
CACCCACGAGGAGCATCGTGGAAAAAGAAGACGTTCCAACCACGTCTTCAAAGCAAGTGGATTGAT
GTGATATCTCCACTGACGTAAGGGATGACGCACAATCCCACTATCCTTCGCAAGACCCTTCCTCTAT
ATAAGGAAGTTCATTTCATTTGGAGAGAACACGGGGGACTCTAGACTGGTACCCGGGTCGATAAGA
GGTTCCAACTTTCACCATAATTAAATAAGATCACTACCGGGCGTATTTTTTGAGTTGTCAATTTTCA[
TG]GAGCTA**AGGA**AGTCGACATGAAAAAGCCTGAACTCACCGCGACGTCTGTCGAGAAGTTTCTGA
TCGAAAAGTTCGACAGCGTCTCCGACCTGATGCAGCTCTCGGAGGGCGAAGAATCTCGTGCTTTCA
GCTTCGATGTAGGAGGGCGTGGATATGTCCTGCGGGTAAATAGCTGCGCCGATGGTTTCTACAAAG
ATCGTTATGTTTATCGGCACTTTGCATCGGCCGCGCTCCCGATTCCGGAAGTGCTTGACATTGGGG
AGTTTAGCGAGAGCCTGACCTATTGCATCTCCCGCCGTGCACAGGGTGTCACGTTGCAAGACCTGC
CTGAAACCGAACTGCCCGCTGTTCTACAACCGGTCGCGGAGGCTATGGATGCGATCGCTGCGGCC
GATCTTAGCCAGACGAGCGGGTTCGGCCCATTCGGACCGCAAGGAATCGGTCAATACACTACATGG
CGTGATTTCATATGCGCGATTGCTGATCCCCATGTGTATCACTGGCAAACTGTGATGGACGACACC
GTCAGTGCGTCCGTCGCGCAGGCTCTCGATGAGCTGATGCTTTGGGCCGAGGACTGCCCCGAAGT
CCGGCACCTCGTGCACGCGGATTTCGGCTCCAACAATGTCCTGACGGACAATGGCCGCATAACAGC
GGTCATTGACTGGAGCGAGGCGATGTTCGGGGATTCCCAATACGAGGTCGCCAACATCTTCTTCTG
GAGGCCGTGGTTGGCTTGTATGGAGCAGCAGACGCGCTACTTCGAGCGGAGGCATCCGGAGCTTG

CAGGATCGCCACGACTCCGGGCGTATATGCTCCGCATTGGTCTTGACCAACTCTATCAGAGCTTGG
TTGACGGCAATTTCGATGATGCAGCTTGGGCGCAGGGTCGATGCGACGCAATCGTCCGATCCGGA
GCCGGGACTGTCGGGCGTACACAAATCGCCCGCAGAAGCGCGGCCGTCTGGACCGATGGCTGTGT
AGAAGTACTCGCCGATAGTGGAAACCGACGCCCCAGCACTCGTCCGAGGGCAAAGAAACTAGTAT
GGTGAGCAAGGGCGAGGAGCTGTTCACCGGGGTGGTGCCCATCCTGGTCGAGCTGGACGGCGAC
GTAAACGGCCACAAGTTCAGCGTGTCCGGCGAGGGCGAGGGCGATGCCACCTACGGCAAGCTGAC
CCTGAAGTTCATCTGCACCACCGGCAAGCTGCCCGTGCCCTGGCCCACCCTCGTGACCACCTTCAC
CTACGGCGTGCAGTGCTTCAGCCGCTACCCCGACCACATGAAGCAGCACGACTTCTTCAAGTCCGC
CATGCCCGAAGGCTACGTCCAGGAGCGCACCATCTTCTTCAAGGACGACGGCAACTACAAGACCCG
CGCCGAGGTGAAGTTCGAGGGCGACACCCTGGTGAACCGCATCGAGCTGAAGGGCATCGACTTCA
AGGAGGACGGCAACATCCTGGGGCACAAGCTGGAGTACAACTACAACAGCCACAACGTCTATATCAT
GGCCGACAAGCAGAAGAACGGCATCAAGGTGAACTTCAAGATCCGCCACAACATCGAGGACGGCAG
CGTGCAGCTCGCCGACCACTACCAGCAGAACACCCCCATCGGCGACGGCCCCGTGCTGCTGCCCGA
CAACCACTACCTGAGCACCCAGTCCGCCCTGAGCAAAGACCCCAACGAGAAGCGCGATCACATGGT
CCTGCTGGAGTTCGTGACCGCCGCCGGGATCACTCACGGCATGGACGAGCTGTACAAGTAAGAGCT
CGAATTTCCCCGATCGTTCAAACATTTGGCAATAAAGTTTCTTAAGATTGAATCCTGTTGCCGGTCT
TGCGATGATTATCATATAATTTCTGTTGAATTACGTTAAGCATGTAATAATTAACATGTAATGCATGA
CGTTATTTATGAGATGGGTTTTTATGATTAGAGTCCCGCAATTATACATTTAATACGCGATAGAAAAC
AAAATATAGCGCGCAAACTAGGATAAATTATCGCGCGCGGTGTCATCTATGTTACTAGATCGGGAAT
TC

P35S-HPT-GFP-TNOS (original version)

35S promoter long enhancer; HPT CDS; GFP CDS; NOS terminator

AGATTAGCCTTTTCAATTTCAGAAAGAATGCTAACCCACAGATGGTTAGAGAGGCTTACGCAGCAGG
TCTCATCAAGACGATCTACCCGAGCAATAATCTCCAGGAAATCAAATACCTTCCCAAGAAGGTTAAA
GATGCAGTCAAAAGATTCAGGACTAACTGCATCAAGAACACAGAGAAAGATATATTTCTCAAGATCA
GAAGTACTATTCCAGTATGGACGATTCAAGGCTTGCTTCACAAACCAAGGCAAGTAATAGAGATTGG
AGTCTCTAAAAAGGTAGTTCCCACTGAATCAAAGGCCATGGAGTCAAAGATTCAAATAGAGGACCTA
ACAGAACTCGCCGTAAAGACTGGCGAACAGTTCATACAGAGTCTCTTACGACTCAATGACAAGAAGA
AAATCTTCGTCAACATGGTGGAGCACGACACACTTGTCTACTCCAAAAATATCAAAGATACAGTCTC
AGAAGACCAAAGGGCAATTGAGACTTTTCAACAAAGGGTAATATCCGGAAACCTCCTCGGATTCCAT
TGCCCAGCTATCTGTCACTTTATTGTGAAGATAGTGGAAAAGGAAGGTGGCTCCTACAAATGCCATC
ATTGCGATAAAGGAAAGGCCATCGTTGAAGATGCCTCTGCCGACAGTGGTCCCAAAGATGGACCCC
CACCCACGAGGAGCATCGTGGAAAAAGAAGACGTTCCAACCACGTCTTCAAAGCAAGTGGATTGAT
GTGATATCTCCACTGACGTAAGGGATGACGCACAATCCCACTATCCTTCGCAAGACCCTTCCTCTAT
ATAAGGAAGTTCATTTCATTTGGAGAGAACACGGGGGACTCTAGACTGGTACCCGGGTCGACATGA
AAAAGCCTGAACTCACCGCGACGTCTGTCGAGAAGTTTCTGATCGAAAAGTTCGACAGCGTCTCCG
ACCTGATGCAGCTCTCGGAGGGCGAAGAATCTCGTGCTTTCAGCTTCGATGTAGGAGGGCGTGGA
TATGTCCTGCGGGTAAATAGCTGCGCCGATGGTTTCTACAAAGATCGTTATGTTTATCGGCACTTT
GCATCGGCCGCGCTCCCGATTCCGGAAGTGCTTGACATTGGGGAGTTTAGCGAGAGCCTGACCTA

TTGCATCTCCCGCCGTGCACAGGGTGTCACGTTGCAAGACCTGCCTGAAACCGAACTGCCCGCTGT
TCTACAACCGGTCGCGGAGGCTATGGATGCGATCGCTGCGGCCGATCTTAGCCAGACGAGCGGGT
TCGGCCCATTCGGACCGCAAGGAATCGGTCAATACACTACATGGCGTGATTTCATATGCGCGATTG
CTGATCCCCATGTGTATCACTGGCAAACTGTGATGGACGACACCGTCAGTGCGTCCGTCGCGCAGG
CTCTCGATGAGCTGATGCTTTGGGCCGAGGACTGCCCCGAAGTCCGGCACCTCGTGCACGCGGAT
TTCGGCTCCAACAATGTCCTGACGGACAATGGCCGCATAACAGCGGTCATTGACTGGAGCGAGGCG
ATGTTCGGGGATTCCCAATACGAGGTCGCCAACATCTTCTTCTGGAGGCCGTGGTTGGCTTGTATG
GAGCAGCAGACGCGCTACTTCGAGCGGAGGCATCCGGAGCTTGCAGGATCGCCACGACTCCGGGC
GTATATGCTCCGCATTGGTCTTGACCAACTCTATCAGAGCTTGGTTGACGGCAATTTCGATGATGC
AGCTTGGGCGCAGGGTCGATGCGACGCAATCGTCCGATCCGGAGCCGGGACTGTCGGGCGTACAC
AAATCGCCCGCAGAAGCGCGGCCGTCTGGACCGATGGCTGTGTAGAAGTACTCGCCGATAGTGGAA
ACCGACGCCCCAGCACTCGTCCGAGGGCAAAGAAAACTAGTATGGTGAGCAAGGGCGAGGAGCTGT
TCACCGGGGTGGTGCCCATCCTGGTCGAGCTGGACGGCGACGTAAACGGCCACAAGTTCAGCGTG
TCCGGCGAGGGCGAGGGCGATGCCACCTACGGCAAGCTGACCCTGAAGTTCATCTGCACCACCGG
CAAGCTGCCCGTGCCCTGGCCCACCCTCGTGACCACCTTCACCTACGGCGTGCAGTGCTTCAGCCG
CTACCCCGACCACATGAAGCAGCACGACTTCTTCAAGTCCGCCATGCCCGAAGGCTACGTCCAGGA
GCGCACCATCTTCTTCAAGGACGACGGCAACTACAAGACCCGCGCCGAGGTGAAGTTCGAGGGCGA
CACCCTGGTGAACCGCATCGAGCTGAAGGGCATCGACTTCAAGGAGGACGGCAACATCCTGGGGC
ACAAGCTGGAGTACAACTACAACAGCCACAACGTCTATATCATGGCCGACAAGCAGAAGAACGGCAT
CAAGGTGAACTTCAAGATCCGCCACAACATCGAGGACGGCAGCGTGCAGCTCGCCGACCACTACCA
GCAGAACACCCCCATCGGCGACGGCCCCGTGCTGCTGCCCGACAACCACTACCTGAGCACCCAGTC
CGCCCTGAGCAAAGACCCCAACGAGAAGCGCGATCACATGGTCCTGCTGGAGTTCGTGACCGCCGC
CGGGATCACTCACGGCATGGACGAGCTGTACAAGTAAGAGCTCGAATTTCCCCGATCGTTCAAACA
TTTGGCAATAAAGTTTCTTAAGATTGAATCCTGTTGCCGGTCTTGCGATGATTATCATATAATTTCT
GTTGAATTACGTTAAGCATGTAATAATTAACATGTAATGCATGACGTTATTTATGAGATGGGTTTTT
ATGATTAGAGTCCCGCAATTATACATTTAATACGCGATAGAAAACAAAATATAGCGCGCAAACTAGGA
TAAATTATCGCGCGCGGTGTCATCTATGTTACTAGATCGGGAATTC

**Supplementary Figure S1. The sequences of P35S-PCAT1-HPT-GFP-TNOS, P35S-PCAT2-HPT-GFP-TNOS, and P35S enhancer-PNOS-HPT-GFP-TNOS.**
